# Iohexol-measured glomerular filtration rate and urinary biomarker changes between vancomycin and vancomycin plus piperacillin-tazobactam in a translational rat model

**DOI:** 10.1101/2023.03.09.532007

**Authors:** Jack Chang, Gwendolyn M. Pais, Sylwia Marianski, Kimberly Valdez, Emily Lesnicki, Erin F. Barreto, Nathaniel J. Rhodes, Paul R. Yarnold, Marc H. Scheetz

## Abstract

Recent clinical studies have reported additive nephrotoxicity with the combination of vancomycin and piperacillin-tazobactam. However, preclinical models have failed to replicate this finding. This study assessed differences in iohexol-measured glomerular filtration rate (GFR) and urinary injury biomarkers among rats receiving this antibiotic combination.

Male Sprague-Dawley rats received either intravenous vancomycin, intraperitoneal piperacillin-tazobactam, or both for 96 hours. Iohexol-measured GFR was used to quantify real-time kidney function changes. Kidney injury was evaluated via the urinary biomarkers: kidney injury molecule-1 (KIM-1), clusterin, and osteopontin.

Compared to the control, rats that received vancomycin had numerically lower GFR after drug dosing on day 3. Rats in this group also had elevations in urinary KIM-1 on experimental days 2 and 4. Increasing urinary KIM-1 was found to correlate with decreasing GFR on experimental days 1 and 3. Rats that received vancomycin+piperacillin-tazobactam did not exhibit worse kidney function or injury biomarkers compared to vancomycin alone.

The combination of vancomycin+piperacillin-tazobactam does not cause additive nephrotoxicity in a translational rat model. Future clinical studies investigating this antibiotic combination should employ more sensitive biomarkers of kidney function and injury, similar to those utilized in this study.

## Introduction

Vancomycin is a glycopeptide antibiotic that remains a treatment of choice for methicillin-resistant *Staphylococcus aureus* and other resistant Gram-positive infections. It is the most frequently prescribed parenteral antibiotic in United States hospitals and utilization has steadily increased in recent years ^1^. Nephrotoxicity is the most common adverse event associated with vancomycin, with conservative estimates attributing rates of >10% ^2–4^. Piperacillin-tazobactam is a broad-spectrum beta-lactam/beta-lactamase inhibitor antibiotic which is commonly prescribed in conjunction with vancomycin for empiric coverage of Gram-negative organisms ^5,6^. Multiple recent clinical meta-analyses have found the combination of vancomycin and piperacillin-tazobactam to be associated with additive nephrotoxicity, as evaluated by increased serum creatinine (SCr), compared to vancomycin alone ^7–11^. In a study including almost 25,000 patients, Luther et al. found the combination of vancomycin+piperacillin-tazobactam to be associated with increased odds of acute kidney injury (AKI) when compared to vancomycin monotherapy (OR: 3.4, 95% CI: 2.57-4.50) or piperacillin-tazobactam monotherapy (OR: 2.7, 95% CI: 1.97-3.69). Notably, this difference in AKI rates persisted when vancomycin+piperacillin-tazobactam was compared to vancomycin plus other comparable beta-lactams (i.e. vancomycin+cefepime or meropenem, OR: 2.68, 95% CI: 1.83-3.91) ^7^. More recently, Bellos et al. obtained similar results in the largest meta-analysis of vancomycin+piperacillin-tazobactam-associated nephrotoxicity. In their study of over 50,000 patients, vancomycin+piperacillin-tazobactam was found to increase the odds of AKI, as assessed by SCr using the Kidney Disease: Improving Global Outcomes criteria, when compared to vancomycin (OR: 2.05, 95% CI: 1.17-3.46), vancomycin+cefepime (OR: 1.80, 95% CI: 1.13-2.77), and vancomycin+meropenem (OR: 1.84, 95% CI: 1.02-3.10) ^10^. Although these results from retrospective studies suggest that vancomycin+piperacillin-tazobactam is associated with greater SCr increases compared to vancomycin alone, it is still unclear whether these changes in SCr reflect loss of kidney function or actual injury. The underlying mechanism of increased SCr associated with vancomycin+piperacillin-tazobactam remains unknown, and SCr is well known to be a poor surrogate marker for both kidney function and injury ^12^.

Despite consistency among the retrospective studies, results may be confounded due to retrospective study design and use of SCr as the singular marker for both kidney injury and function. SCr is not a direct marker for kidney function; rather, it is an imperfect surrogate for glomerular filtration rate (GFR). Changes in the secretion and/or reabsorption of creatinine can result in SCr fluctuations that are not reflective of true kidney function decline ^12–14^. Additionally, SCr is a poor surrogate for identification of acute kidney injury; in the acute setting, GFR can decline by as much as 50% prior to detectable rises in SCr ^14^. Clinically, this delay in detectable SCr rise impedes the early detection of AKI and corresponding dose adjustments to renally-eliminated medications.

In order to develop more sensitive and specific biomarkers for kidney injury, several translational animal studies have investigated the injury profile of vancomycin+piperacillin-tazobactam vs. vancomycin using newer biomarkers of kidney injury (e.g. kidney injury molecule-1 [KIM-1], clusterin, and osteopontin [OPN]) and histopathological analysis ^15,16^. In contrast to retrospective human studies, these animal models showed that the combination of vancomycin+piperacillin-tazobactam did not result in excess kidney injury. In fact, co-administration of piperacillin-tazobactam may be nephroprotective during vancomycin administration ^15–17^. Animal studies markedly differed from human studies in that they utilized sensitive and specific kidney injury biomarkers such as KIM-1, clusterin, and OPN in the detection of antibiotic-induced nephrotoxicity ^18^. KIM-1 has been identified as the most relevant urinary biomarker in animal models, correlating with both early GFR changes and end-organ histopathologic damage at the proximal tubule ^15,18–20^.

Newer markers of kidney function are also under investigation. Traditionally, inulin has been a gold-standard marker for GFR estimation. However, practical use is limited by the extensive labor and high costs associated with implementation ^21–23^. Other alternative markers such as radiolabeled tracers (e.g. ethylenediaminetetraacetic acid, triamine pentaacetic acid, ^51^Cr-EDTA, ^99m^Tc-DTPA, and ^125^I-iothalamte) have shown comparable accuracy and reliability in determining GFR ^24–26^. However, these compounds require special handling, storage, and disposal which complicates use. Iohexol is a low-cost, non-ionic contrast agent that fulfills the criteria for ideal kidney function biomarkers. It is freely filtered and eliminated by the kidneys, non-secreted, non-reabsorbed, and non-metabolized, with low plasma protein binding ^27,28^. There is also significant clinical experience which supports the safety and utility of iohexol ^29–32^.

Recently, we utilized the same experimental design in a translational rat model to investigate the injury profile of vancomycin+piperacillin-tazobactam with an alternative glomerular function biomarker, fluorescein isothiocyanate conjugated sinistrin ^16^. Our findings were concordant from prevoius preclinical studies of this antibiotic combination, in that kidney function changes were observed among rats that received vancomycin alone, but not among those that received vancomycin+piperacillin-tazobactam. Similarly, the injury biomarker, kidney injury molecule-1, was significantly elevated only among rats that received vancomycin alone.

In the present study, we employed our translational rat model to assess kidney function by iohexol clearance as a surrogate for GFR, and kidney injury by urinary biomarkers in rats receiving vancomycin +/− piperacillin-tazobactam, piperacillin-tazobactam alone, or control (saline). The purpose of this study was to directly assess kidney function between treatment groups by GFR, and to correlate GFR to urinary injury biomarkers in rats receiving vancomycin +/− piperacillin-tazobactam, piperacillin-tazobactam alone, or control (saline).

## Results

### Characteristics of animal cohort

A total of 24 male Sprague Dawley rats were studied, with animal dosing group assignments shown in Supplementary figure 1. For the pharmacokinetic analysis, two animals provided only terminal plasma samples due to occluded catheters; however, individual Bayesian posteriors were generated for all animals. All other animals contributed complete pharmacokinetic data to the model, and all animals were included in the pharmacodynamic analyses. Because of a sample collection error, baseline (day 0) urine samples were not collected for rats assigned to the vancomycin+piperacillin-tazobactam treatment group.

### Model characteristics

To describe plasma iohexol clearance, a two-compartment intravenous infusion base model with first-order elimination fit the data well (Supplementary figure 2). The final model’s population mean parameter values (SD) for clearance (CL) and central compartment volume (V1), were 0.076 L/hr (0.0027) and 0.03 L (0.0023), respectively. Peripheral compartment volume (V2), and intercompartmental transfer (Q) were fixed at 0.0067 L and 0.018 L/hr, respectively. The linear regression of the observed concentrations versus Bayesian posterior population predictions resulted in an intercept of 1.34, with a slope of 0.99 and *R*^2^ value of 0.92. The linear regression of the observed concentrations versus Bayesian posterior individual predictions resulted in an intercept of 1.01 with a slope of 1.02 and *R*^2^ value of 0.97 (Supplementary figure 3).

### Kidney function over time

Compared to the control group, rats that received only vancomycin had a lower GFR on day 3 (Figure 1: -0.04 mL/min/100g body weight; 95% confidence interval [CI]: -0.09 to -0.01; p<0.05). No other significant changes in GFR were identified, including rats in the vancomycin+piperacillin-tazobactam group. Baseline GFR was not assessed in this protocol since plasma samples were not obtained at baseline (day 0). Baseline creatinine clearance (CrCL) was not different between the vancomycin, piperacillin-tazobactam, and control groups [with control used as the referent group] (0.51 ± 0.13, 0.52 ± 0.12, and 0.52 ± 0.09 mL/min/100 g body weight; p = 0.98). Baseline CrCL was not able to be assessed for the vancomycin+piperacillin-tazobactam group rats. Following administration of the first dose (day 1), rats in the vancomycin+piperacillin-tazobactam group experienced a slight decline in CrCL which was not seen among the other treatment groups. No differences in CrCL were identified in the other treatment groups (Figure 2).

**Figure 1:**
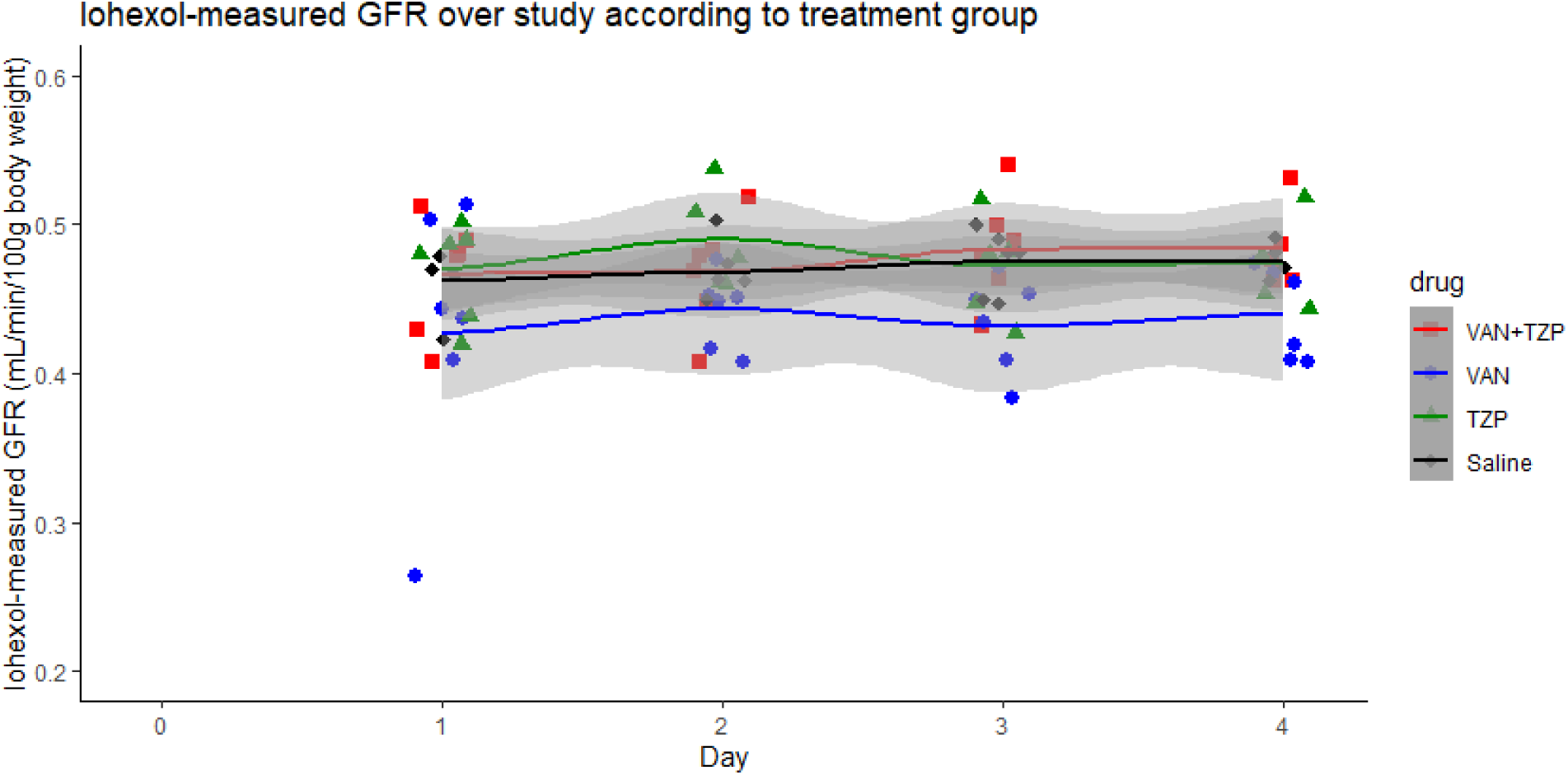
Comparison of individual iohexol glomerular filtration rate (GFR) measurements between treatment groups and across dosing days. Compared to the control group, rats in the VAN (vancomycin) group [blue] showed a decline in GFR over dosing days 1-4, with a significant difference seen on day 3 (-0.04 mL/min/100g body weight; 95% confidence interval [CI]: -0.09 to -0.01; p=0.046). Colored lines depict the locally weighted scatterplot smoothing (LOWESS) trendline for each respective treatment groups; 95% confidence intervals for LOWESS trendlines are depicted by the gray shaded areas.

**Figure 2:**
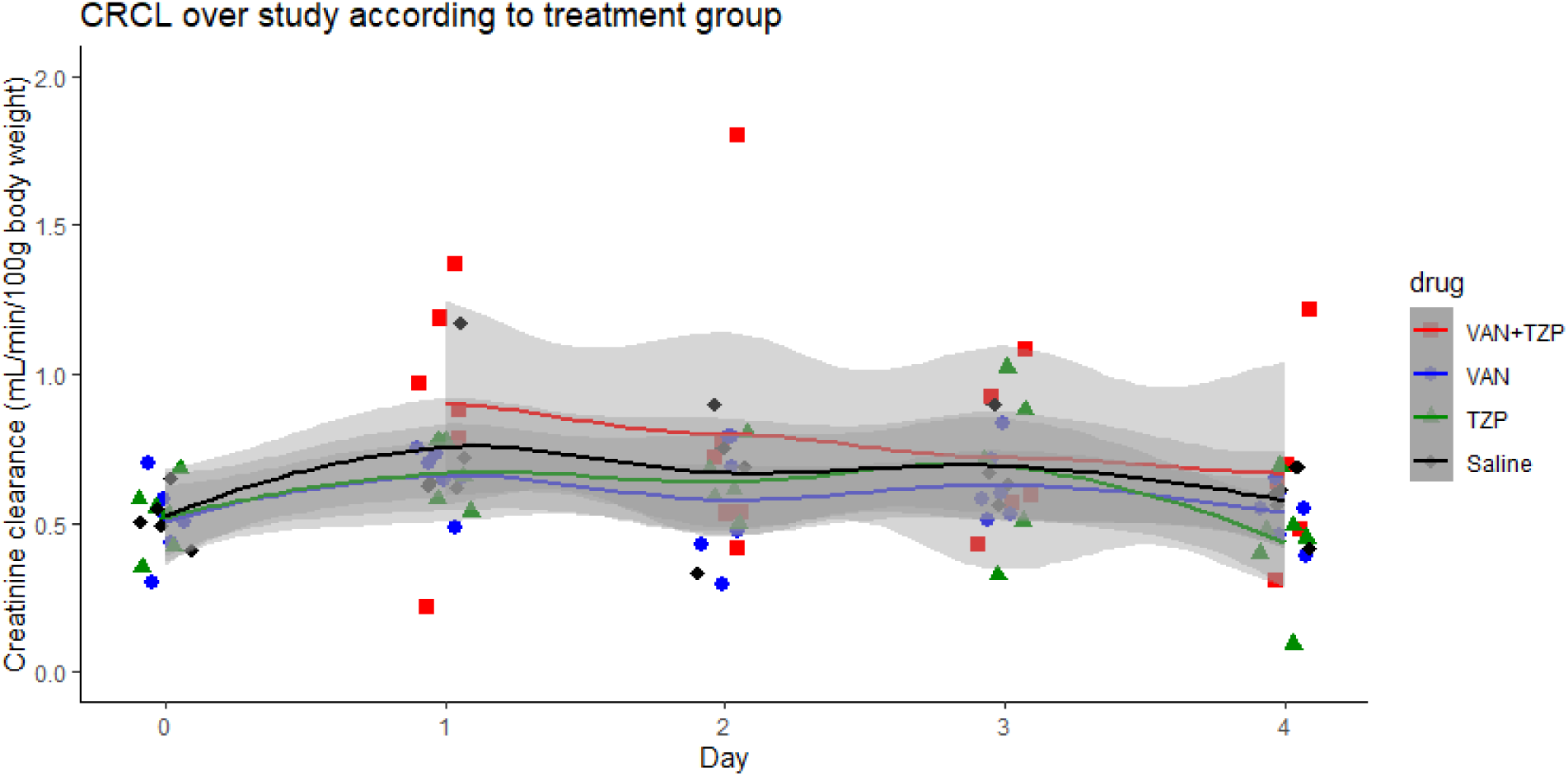
Comparison of individual creatinine clearance (CrCL) measurements between treatment groups and across dosing days. Using the control group as the referent, no significant differences in CrCL were identified. Colored lines depict the LOWESS trendline for each respective treatment groups; 95% confidence intervals for LOWESS trendlines are depicted by the gray shaded areas.

### Urinary output and injury biomarkers

Summary data for urinary output and injury biomarker data are listed in Table 1. Baseline urinary output was not different among the treatment groups. When daily urine output among the treatment groups was compared to the control, no significant differences were observed.

**Table 1:** Summary of weight loss, urinary output, and urinary biomarkers.

| Characteristic | Treatment group |  |  |  |
| --- | --- | --- | --- | --- |
|  | Vancomycin | Piperacillin-tazobactam | Vancomycin+piperacillin-tazobactam | Control |
| Number of animals | 6 | 6 | 6 | 6 |
| Baseline weight (g), mean ± SD | 271.9 ± 11.1 | 282.0 ± 16.9 | 274.6 ± 11.6 | 275.2 ± 8.9 |
| Weight change (g), mean ± SD | +3.3 ± 10.0 | +3.8 ± 7.3 | -0.3 ± 15.4 | +0.02 ± 10.8 |
| Urine output (mL), mean ± SD | | | | |
| Baseline | 5.6 ± 3.2 | 6.8 ± 2.9 | --- | 7.5 ± 7.2 |
| Day 1 | 6.6 ± 3.2 | 8.1 ± 2.3 | 6.2 ± 3.2 | 9.8 ± 5.1 |
| Day 2 | 6.3 ± 2.9 | 7.0 ± 2.9 | 5.8 ± 2.5 | 8.4 ± 1.9 |
| Day 3 | 7.7 ± 2.5 | 8.1 ± 4.2 | 5.8 ± 3.4 | 8.5 ± 4.3 |
| Day 4 | 8.6 ± 2.2 | 8.3 ± 3.1 | 6.4 ± 3.5 | 8.7 ± 1.0 |
| KIM-1 (ng), median (IQR) |  |  |  |  |
| Baseline | 12.1 (11.3 to 14.1) | 7.76 (4.88 to 10.9) | -- | 6.89 (5.68 to 12.8) |
| Day 1 | 14.1 (10.44 to 16.56) | 6.50 (5.48 to 7.70) | 13.6 (10.7 to 14.5) | 7.72 (7.20 to 7.84) |
| Day 2 | 22.6 (8.50 to 35.1) | 6.05 (5.60 to 7.08) | 12.3 (8.36 to 15.5) | 8.68 (7.30 to 8.80) |
| Day 3 | 20.5 (16.4 to 39.1) | 6.23 (4.64 to 7.42) | 11.9 (8.56 to 22.3) | 8.33 (7.42 to 9.58) |
| Day 4 | 28.1 (16.1 to 41.3) | 5.74 (5.26 to 7.58) | 13.8 (9.92 to 20.9) | 7.84 (7.68 to 8.94) |
| Clusterin (ng), median (IQR) |  |  |  |  |
| Baseline | 3956 (2974 to 6859) | 2729 (1945 to 5596) | -- | 2536 (1830 to 5299) |
| Day 1 | 4053 (3177 to 5698) | 2883 (1807 to 4093) | 6603 (4824 to 8713) | 3222 (2725 to 4273) |
| Day 2 | 6177 (2328 to 6881) | 2998 (1750 to 3885) | 5324 (3382 to 7176) | 2496 (2444 to 2977) |
| Day 3 | 6481 (5707 to 11468) | 2311 (1137 to 3161) | 4788 (3232 to 5533) | 2793 (2364 to 3292) |
| Day 4 | 6902 (5106 to 9429) | 3213 (2468 to 3630) | 5062 (4147 to 6328) | 2480 (2444 to 2715) |
| Osteopontin (ng), median (IQR) |  |  |  |  |
| Baseline | 1.91 (1.17 to 2.63) | 0.92 (0.56 to 1.39) | -- | 0.68 (0.48 to 0.80) |
| Day 1 | 3.56 (2.16 to 4.64) | 1.21 (1.08 to 1.28) | 1.95 (1.36 to 3.36) | 0.81 (0.40 to 1.18) |
| Day 2 | 3.22 (1.92 to 4.32) | 1.27 (0.71 to 1.38) | 2.44 (1.76 to 3.36) | 0.81 (0.68 to 0.96) |
| Day 3 | 3.63 (2.80 to 4.48) | 0.98 (0.64 to 1.43) | 1.95 (1.51 to 3.25) | 0.60 (0.46 to 0.88) |
| Day 4 | 4.16 (3.39 to 4.83) | 1.30 (0.63 to 1.53) | 2.88 (2.21 to 3.50) | 0.52 (0.49 to 0.53) |
| GFR (mL/min/100 g B.W.), mean ± SD | | | | |
| Day 1 | 0.43 ± 0.09 | 0.47 ± 0.03 | 0.47 ± 0.04 | 0.46 ± 0.02 |
| Day 2 | 0.44 ± 0.03 | 0.49 ± 0.03 | 0.47 ± 0.04 | 0.47 ± 0.02 |
| Day 3 | 0.43 ± 0.03 | 0.47 ± 0.03 | 0.48 ± 0.04 | 0.48 ± 0.02 |
| Day 4 | 0.44 ± 0.03 | 0.48 ± 0.03 | 0.48 ± 0.03 | 0.48 ± 0.01 |
| CrCL (mL/min/100 g B.W.), mean ± SD | | | | |
| Baseline | 0.51 ± 0.13 | 0.52 ± 0.12 | -- | 0.52 ± 0.09 |
| Day 1 | 0.66 ± 0.09 | 0.67 ± 0.11 | 0.83 ± 0.33 | 0.75 ± 0.24 |
| Day 2 | 0.58 ± 0.21 | 0.64 ± 0.11 | 0.71 ± 0.30 | 0.67 ± 0.24 |
| Day 3 | 0.63 ± 0.12 | 0.69 ± 0.28 | 0.72 ± 0.27 | 0.69 ± 0.15 |
| Day 4 | 0.53 ± 0.09 | 0.43 ± 0.19 | 0.67 ± 0.34 | 0.55 ± 0.21 |
| Histopathology scores, median (IQR) |  |  |  |  |
| Worst overall score | 1 (1 to 2) | 1 (1 to 1) | 1 (1 to 1) | 1 (1 to 1) |
| Worst tubular score | 1 (1 to 2) | 1 (1 to 1) | 1 (1 to 1) | 1 (1 to 1) |
| Composite tubular score | 3.5 (3 to 8) | 2.5 (2 to 4) | 1.5 (1 to 2) | 3 (2 to 4) |

When each of the treatment groups was compared to the control group, rats in the vancomycin group had the highest urinary KIM-1, with differences seen after drug dosing on day 2 (Figure 3, 33.1 ng, 95% CI: 5.35 to 60.9, p=0.019), and day 4 (38.8 ng, 95% CI: 9.89 to 67.7, p=0.009**). Similar to KIM-1, urinary clusterin was higher only among rats in the vancomycin group on day 2 (Figure 4, 8580 ng, 95% CI: 2682 to 14478, p=0.004**) and day 3 (7108 ng, 95% CI: 1210 to 13005, p=0.011**). Urinary OPN was higher among vancomycin group rats on day 1 (Figure 5, 2.54 ng, 95% CI: 1.54 to 3.56, p<0.001), day 2 (2.28 ng, 95% CI: 1.27 to 3.28, p<0.001), day 3 (3.02 ng, 95% CI: 2.01 to 4.03, p<0.001**), and day 4 (3.54 ng, 95% CI: 2.50 to 4.58, p<0.001**). Urinary OPN was also elevated among vancomycin+piperacillin-tazobactam group rats on day 1 (1.31 ng, 95% CI: 0.29 to 2.31, p=0.011), day 2 (1.77 ng, 95% CI: 0.76 to 2.77, p=0.001), day 3 (1.71 ng, 95% CI: 0.69 to 2.71, p=0.001), and day 4 (2.46 ng, 95% CI: 1.42 to 3.51, p<0.001). Overall, the biomarkers of proximal tubule injury KIM-1 and clusterin, were elevated by day 2 among rats that received vancomycin only. OPN, which is more of a general marker of damage throughout the nephrons, was elevated in both vancomycin and vancomycin+piperacillin-tazobactam groups on days 1-4. In the highly conservative Bonferroni corrections for multiple comparisons, few comparisons remained significant. Osteopontin was significantly different between saline and vancomycin on days 1-4 and vancomycin +piperacillin-tazobactam on days 2- 4 (P<0.05 corrected). No comparisons remained significant when comparing all treatments to controls and adjusted for every possible comparison over time for KIM-1 and clusterin.

**Figure 3:**
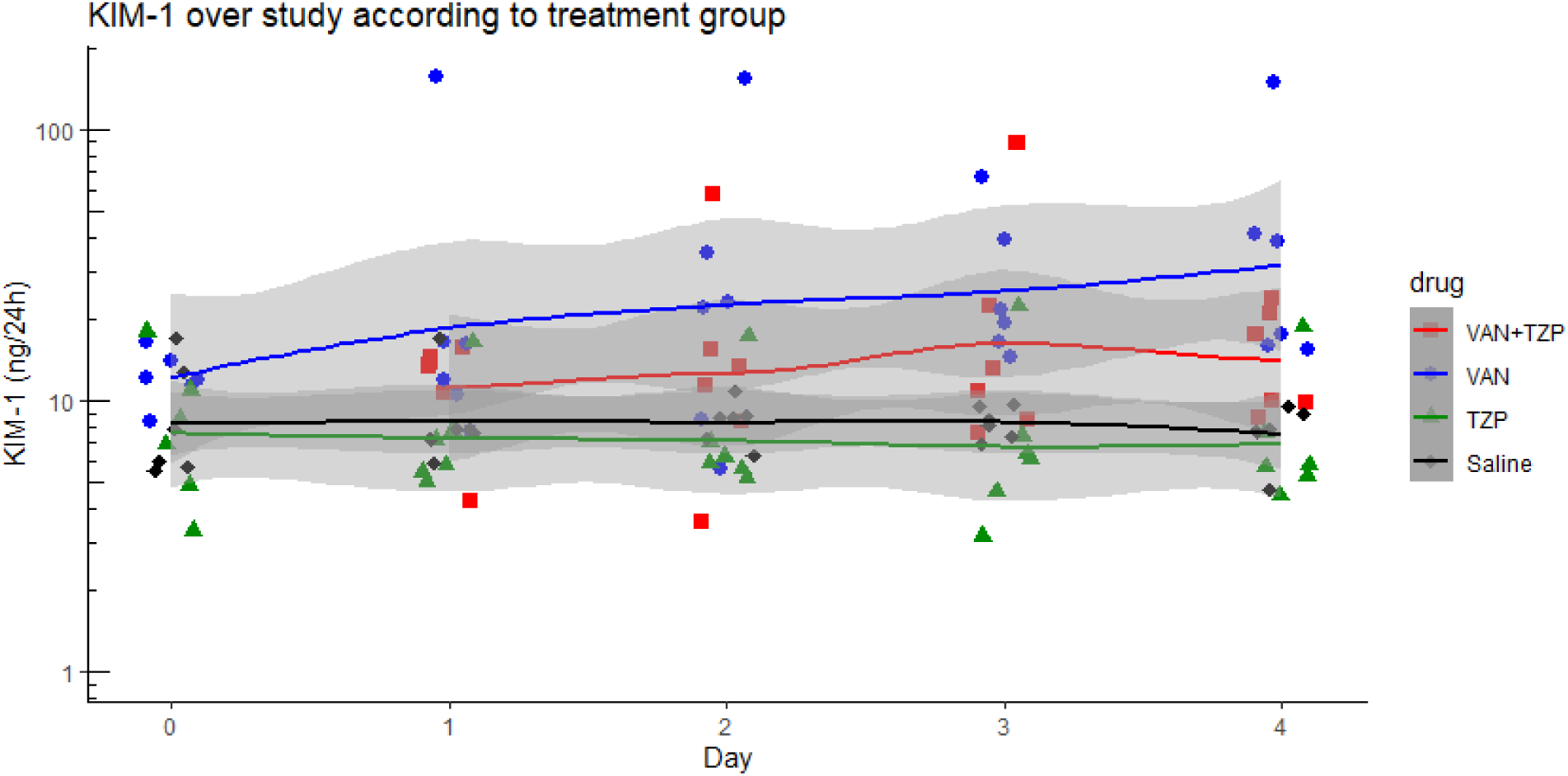
Comparison of individual daily urinary kidney injury molecule-1 (KIM-1) measurements between treatment groups and across dosing days. Using the control as the referent group, rats in the vancomycin group had the highest urinary KIM-1, with significant differences seen after drug dosing on day 2 (33.1 ng, 95% CI: 5.35 to 60.9, p=0.019), and day 4 (38.8 ng, 95% CI: 9.89 to 67.7, p=0.009). Colored lines depict the LOWESS trendline for each respective treatment groups; 95% confidence intervals for LOWESS trendlines are depicted by the gray shaded areas.

**Figure 4:**
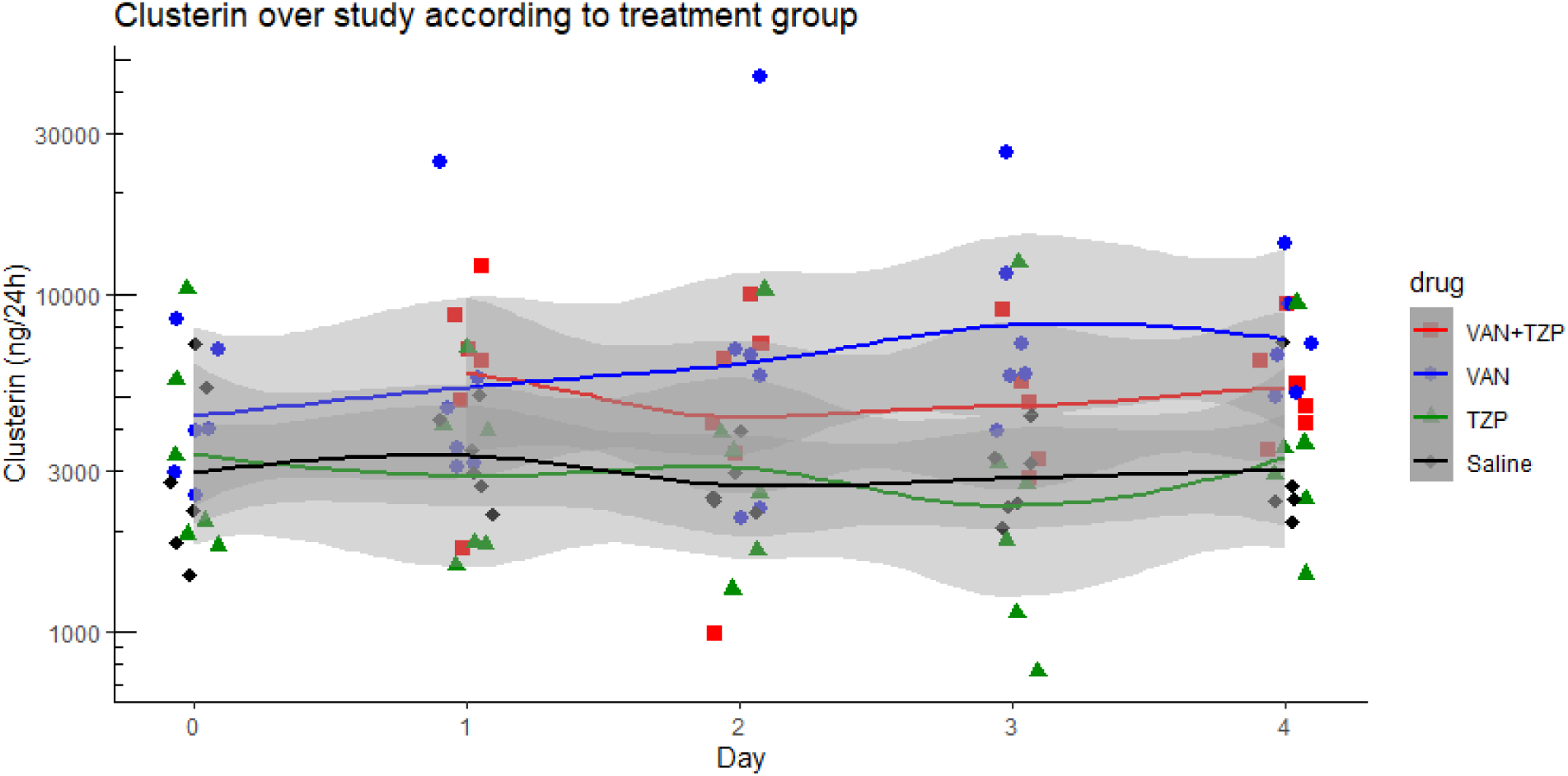
Comparison of individual daily urinary clusterin measurements between treatment groups and across dosing days. Using the control as the referent group, urinary clusterin was significantly higher among rats in the vancomycin group on day 2 (8580, 95% CI: 2682 to 14478, p=0.004) and day 3 (7108, 95% CI: 1210 to 13005, p=0.011). Colored lines depict the LOWESS trendline for each respective treatment groups; 95% confidence intervals for LOWESS trendlines are depicted by the gray shaded areas.

**Figure 5:**
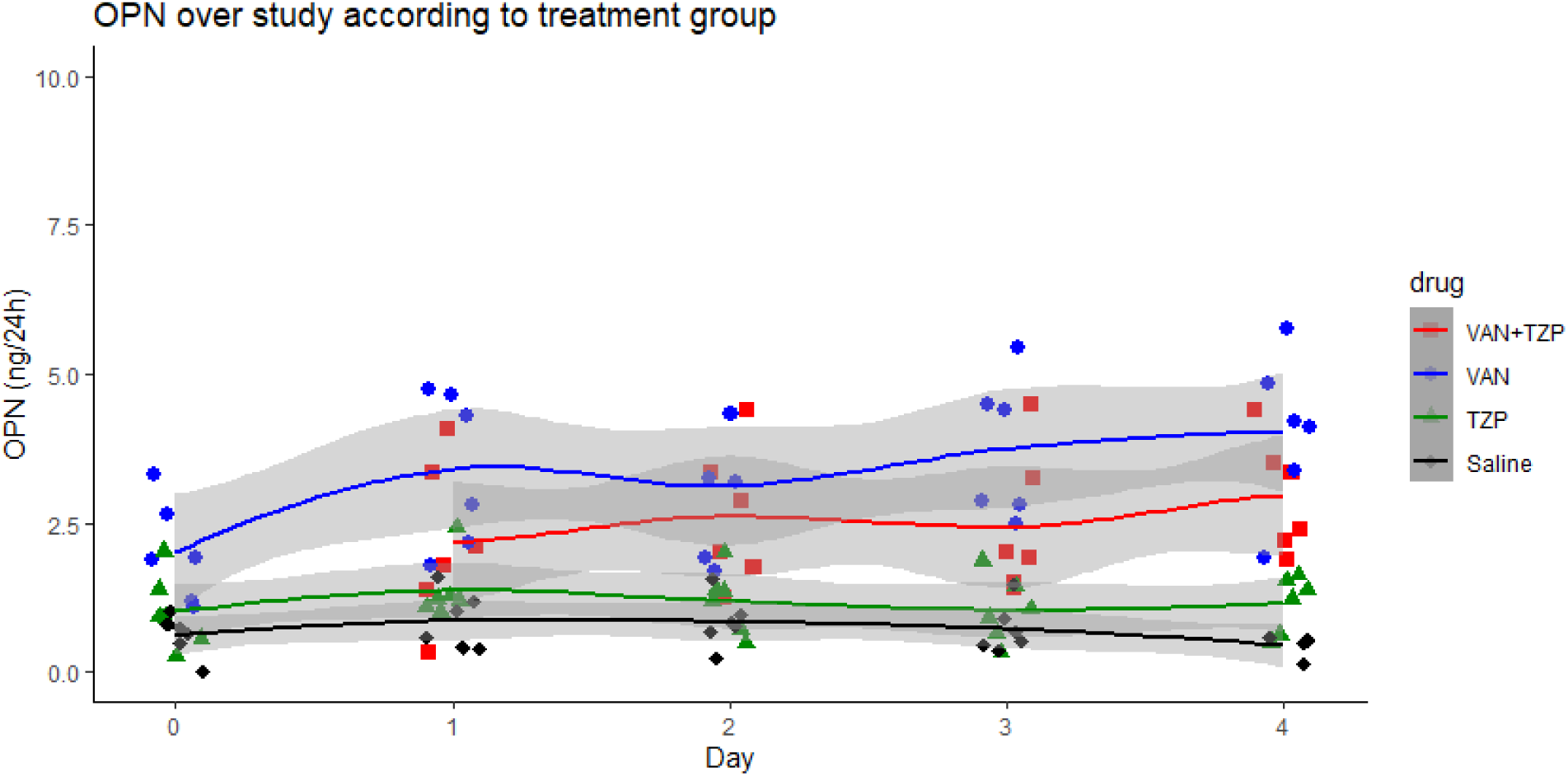
Comparison of individual daily urinary osteopontin (OPN) measurements between treatment groups and across dosing days. Using the control as the referent group, urinary OPN was significantly higher among vancomycin group rats on day 1 (2.55, 95% CI: 1.54 to 3.56, p<0.001), day 2 (2.28, 95% CI: 1.27 to 3.28, p<0.001), day 3 (3.02, 95% CI: 2.01 to 4.03, p<0.001), and day 4 (3.54, 95% CI: 2.50 to 4.58, p<0.001). Urinary OPN was also significantly elevated among vancomycin+piperacillin-tazobactam group rats on day 1 (1.31, 95% CI: 0.29 to 2.31, p=0.011), day 2 (1.77, 95% CI: 0.76 to 2.77, p=0.001), day 3 (1.71, 95% CI: 0.69 to 2.71, p=0.001), and day 4 (2.46, 95% CI: 1.42 to 3.51, p<0.001).

### Correlation between urinary biomarkers of injury and GFR

Spearman’s rank correlation between the urinary biomarkers and GFR measurements are listed in Table 2. Among the vancomycin group rats, urinary KIM-1 was significantly correlated with decreasing GFR on day 1 (Figure 6, Spearman’s rho, -0.43, p=0.037) and day 3 (Spearman’s rho, -0.43, p=0.042). No significant correlations between GFR and clusterin or OPN were identified.

**Figure 6:**
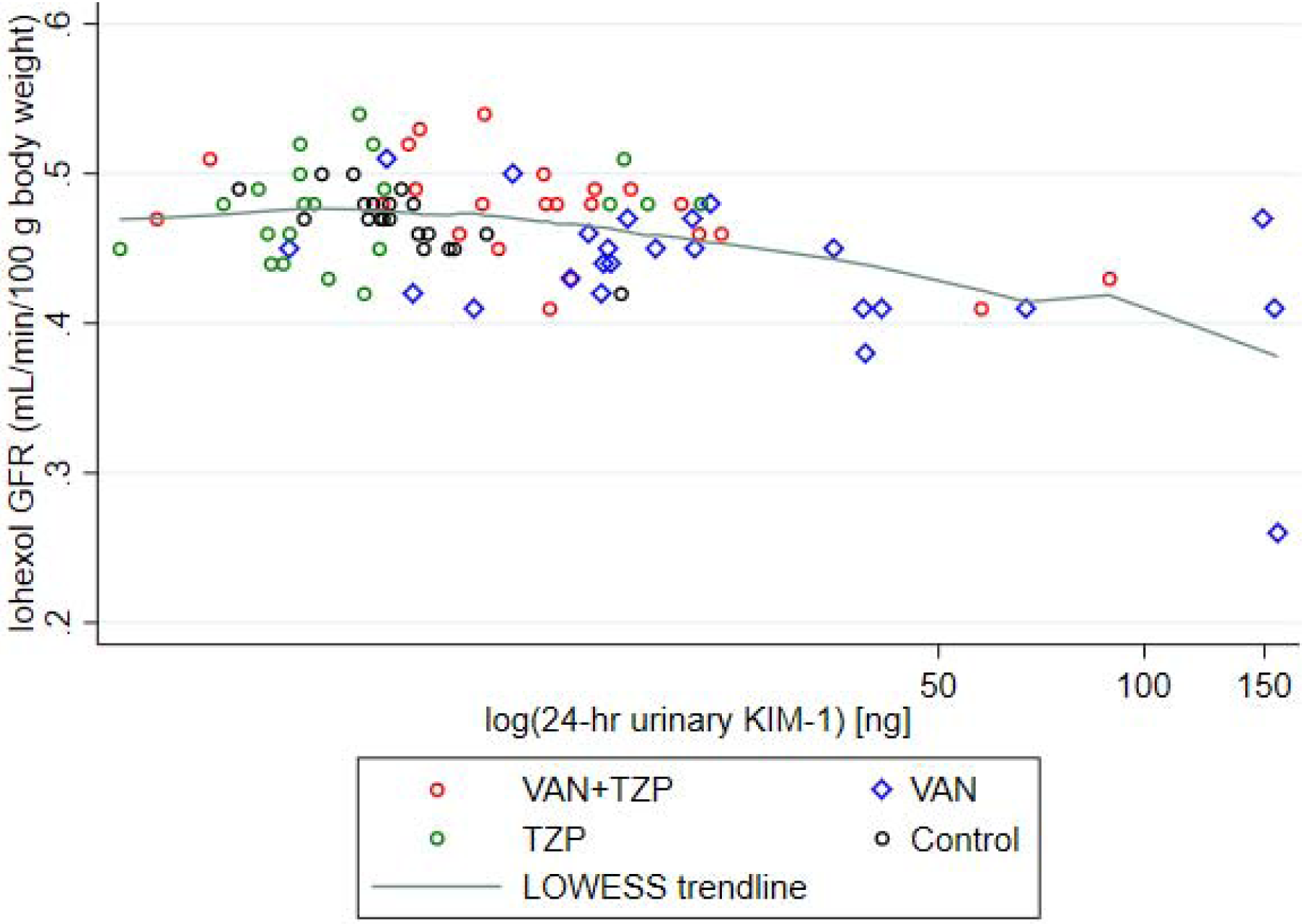
Spearman correlation of GFR with log-normalized, 24-hr urinary KIM-1. Among rats that received vancomycin, urinary KIM-1 was significantly correlated with decreasing GFR on day 1 (Spearman’s rho, -0.43, p=0.037) and day 3 (Spearman’s rho, -0.43, p=0.042). Individual rats are depicted by each data point and the green line depicts the LOWESS trendline.

**Table 2:**
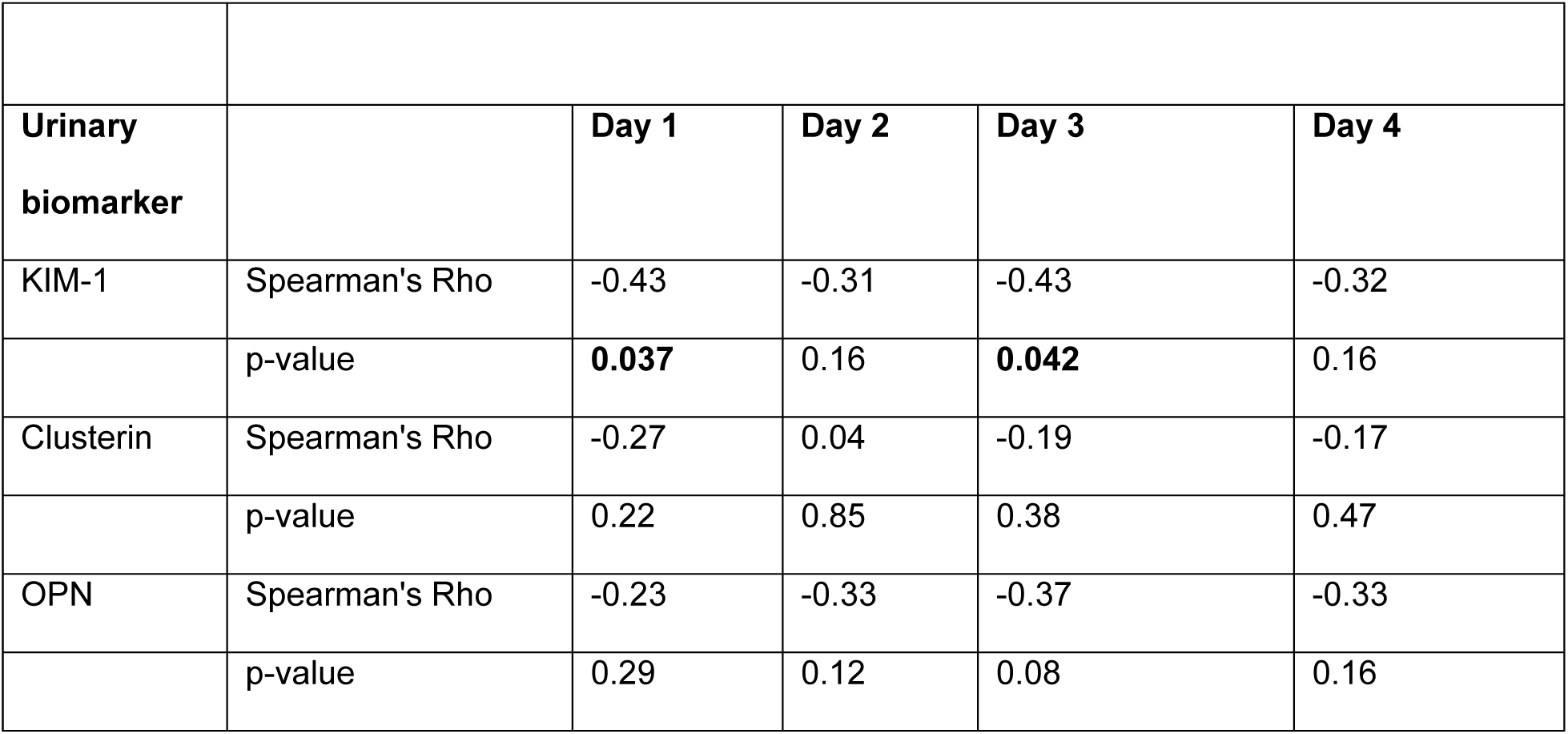
Summary of urinary biomarker correlations with GFR.

|  | **<u>Table 2: Summary of urinary biomarker correlations with GFR</u>** |  |  |  |  |
| --- | --- | --- | --- | --- | --- |
| **Urinary biomarker** |  | **Day 1** | **Day 2** | **Day 3** | **Day 4** |
| KIM-1 | Spearman's Rho | -0.43 | -0.31 | -0.43 | -0.32 |
|  | p-value | **0.037** | 0.16 | **0.042** | 0.16 |
| Clusterin | Spearman's Rho | -0.27 | 0.04 | -0.19 | -0.17 |
|  | p-value | 0.22 | 0.85 | 0.38 | 0.47 |
| OPN | Spearman's Rho | -0.23 | -0.33 | -0.37 | -0.33 |
|  | p-value | 0.29 | 0.12 | 0.08 | 0.16 |

### Correlation between urinary biomarkers of injury and CrCL

Spearman’s rank correlation between the urinary biomarkers and CrCL measurements are listed in Table 3. Among rats that received vancomycin+piperacillin-tazobactam and vancomycin alone, urinary clusterin was significantly correlated with increasing CrCL on day 1 (Figure 7, Spearman’s rho, 0.73, p<0.001), day 2 (Spearman’s rho, 0.63, p<0.003), and day 3 (Spearman’s rho, 0.46, p=0.047). No significant correlations were identified between CrCL with KIM-1 and OPN.

**Figure 7:**
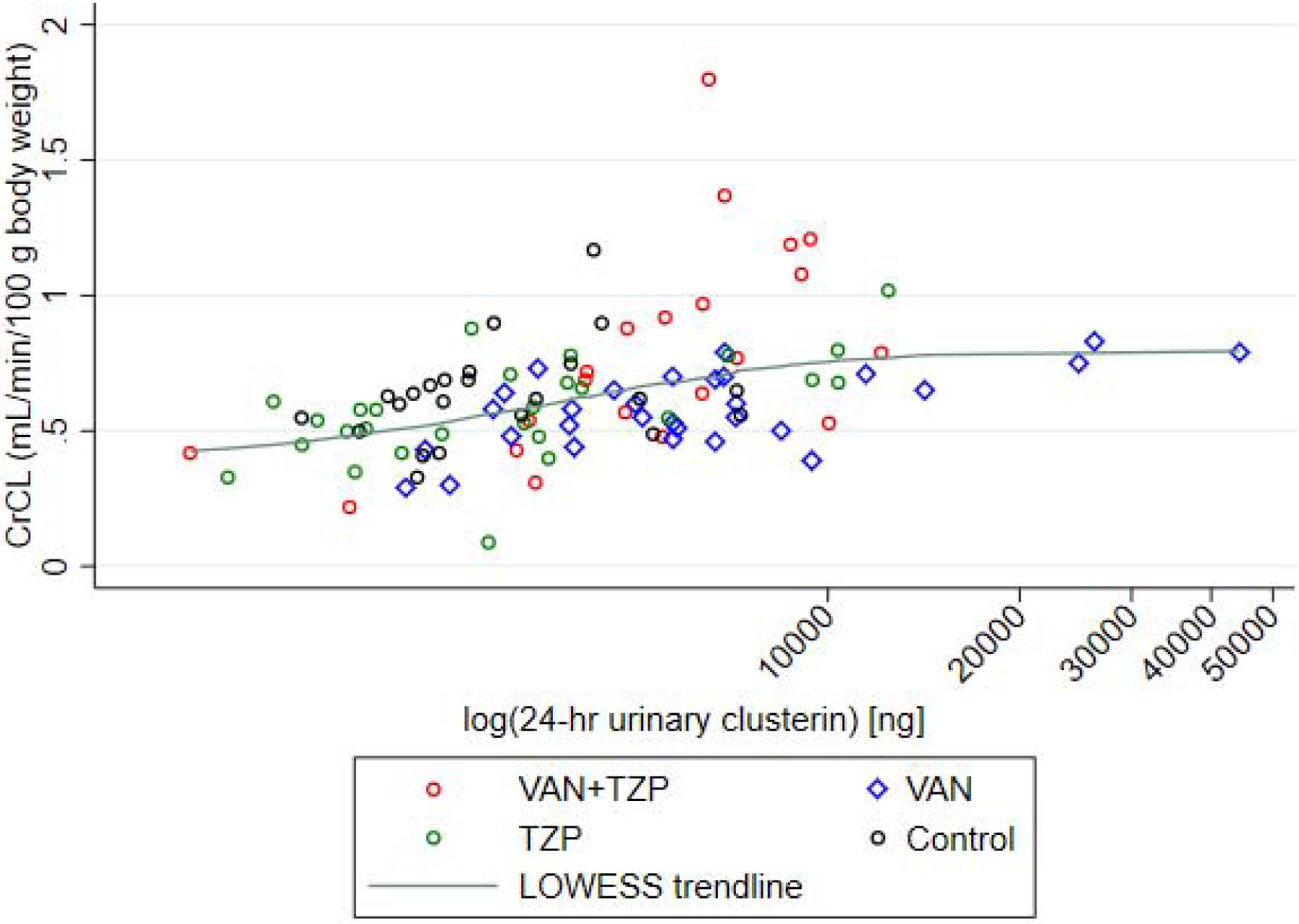
Spearman correlation of CrCL with 24-hr urinary clusterin. Among rats that received vancomycin+piperacillin-tazobactam, urinary clusterin was significantly correlated with increasing CrCL on day 1 (Spearman’s rho, 0.73, p<0.001), day 2 (Spearman’s rho, 0.63, p<0.003), and day 3 (Spearman’s rho, 0.46, p=0.047). Individual rats are depicted by each data point and the green line depicts the LOWESS trendline.

**Table 3:** Summary of urinary biomarker correlations with CrCL.

|  | **<u>Table 3: Summary of urinary biomarker correlations with CrCL</u>** |  |  |  |  |  |
| --- | --- | --- | --- | --- | --- | --- |
| **Urinary biomarker** |  | **Baseline** | **Day 1** | **Day 2** | **Day 3** | **Day 4** |
| KIM-1 | Spearman's Rho | 0.47 | 0.35 | 0.42 | 0.28 | 0.33 |
|  | p-value | 0.06 | 0.11 | 0.06 | 0.23 | 0.14 |
| Clusterin | Spearman's Rho | 0.44 | 0.73 | 0.63 | 0.46 | 0.31 |
|  | p-value | 0.08 | **0.001** | **<0.003** | **0.047** | 0.16 |
| OPN | Spearman's Rho | 0.20 | 0.22 | 0.11 | -0.15 | -0.03 |
|  | p-value | 0.44 | 0.32 | 0.64 | 0.52 | 0.91 |

### Histopathological analysis

Kidney histopathology was variable (Table 1). The vancomycin alone group demonstrated the highest PSTC scores for composite tubular damage (median 3.5; interquartile range [IQR] 3 to 8). Signs of tubular cell alterations consisting of cellular sloughing and granular intratubular casts were only observed among rats that received vancomycin alone. Interstitial inflammation of the cortex was also markedly higher among rats that received vancomycin alone. These indicators of tubular epithelial damage were observed at lower frequency and severity in kidneys from rats that received vancomycin+piperacillin-tazobactam.

## Discussion

This study analyzed GFR, CrCL, urinary injury biomarkers, and histopathology in rats after they received vancomycin, vancomycin+piperacillin-tazobactam, piperacillin-tazobactam, or saline. We found only a minor numeric difference in GFR (as measured by iohexol clearance) among rats that received only vancomycin on day 3 of dosing, when compared to controls that received saline. Elevations in urinary KIM-1 and clusterin, as markers of proximal tubule injury, were evident as early as day 2 (Supplementary figure 4 & 5). KIM-1 correlated with decreasing GFR on days 1 and 3 among rats that received only vancomycin. Osteopontin, a more general marker of damage throughout the nephrons, was elevated among both vancomycin+piperacillin-tazobactam and vancomycin group rats on days 1-4 (Supplementary figure 6). Observed elevations in the injury biomarkers KIM-1 and clusterin are concordant with previous animal studies, which have found these biomarkers to be sensitive and specific for vancomycin-induced kidney damage ^18^.

Notably, vancomycin+piperacillin-tazobactam treated animals did not fare worse than vancomycin only treated animals; in fact, rats that received vancomycin+piperacillin-tazobactam had no significant changes in either GFR or urinary KIM-1 throughout the experiment. Our results are concordant with previous animal studies investigating the injury biomarker KIM-1, in that renal injury was seen earlier and to a greater extent in the vancomycin alone group compared to the vancomycin+piperacillin-tazobactam group ^15,16,19,20^.

Previously, we demonstrated that small changes in GFR (as assessed by fluorescein isothiocyanate-labelled sinistrin [FITC-sinistrin] clearance) were detectable by day 4 in rats that received the same experimental treatment ^16^. In the present study, the magnitude of iohexol GFR change from day 1 to day 4 among all treatment groups (Supplementary figure 7, mean GFR change: 0.01 mL/min/100 g body weight, standard deviation [SD]: 0.073) was numerically lower than GFR changes as assessed by FITC-sinistrin clearance (mean GFR change: 0.057 mL/min/100 g body weight, SD: 0.395) ^16^. Analysis of GFR changes among rats that received only vancomycin in this study showed no change in iohexol GFR from day 1 (mean GFR change: 0.01 mL/min/100 g body weight, SD: 0.113); whereas our previous experiment utilizing FITC-sinistrin for measurement of GFR found a more profound decline in kidney function among rats that received only vancomycin (mean GFR change: -0.39 mL/min/100 g body weight, SD: 0.139). Although the changes in observed GFR among rats that received only vancomycin in this study were smaller than previous studies, the identified GFR drop on day 3 among rats that received only vancomycin still represented a ∼10% decline in GFR when compared to the control group on day 3. The results of the present study are consistent with previous study findings that the combination of vancomycin+piperacillin-tazobactam was not associated with significant declines in GFR when compared to rats that received vancomycin only or controls.

In the present study, we did not identify any changes in kidney function by creatinine clearance (Supplementary figure 8). In clinical studies, CrCL changes are usually seen beyond 3 days There are several possible explanations for differences in our study: 1) CrCL changes may be differ in rats; however, in our study iohexol-measured GFR was generally unchanged over this same time period. 2) clinically observed decline in CrCL may represent ‘pseudonephrotoxicity’ caused by vancomycin+piperacillin-tazobactam. It should be considered that serum creatinine is an endogenous compound, and thus subject to more potential inter- and intra-individual variability including secretion, compared to an exogenous compound such as iohexol which is more instructive for estimation of the true GFR.. It is notable that we did not see a divergence between clearance estimated by iohexol or creatinine in our model. Almost all trends for worsened kidney function and injury were in the vancomycin group versus the vancomycin + piperacillin-tazobactam group. Thus, the results presented here are confirmatory with previous animal models; no preclinical model has been able to recapitulate worse kidney injury or function with vancomycin + piperacillin tazobactam.

Histopathological analysis results are also consistent with our previous findings that higher urinary KIM-1 coaligned with worse histopathological damage scores in the vancomycin-treated rat ^15,16,19^. Clusterin and OPN are more general biomarkers of nonspecific tubule or glomerular damage, with clusterin having been shown to be an earlier predictor of kidney injury compared to OPN ^18^. We found elevations in urinary clusterin on day 3, among rats that received vancomycin+piperacillin-tazobactam. A significant correlation between increasing clusterin and increasing CrCL was also identified on day 1 and day 2, among rats that received vancomycin+piperacillin-tazobactam. It is less clear what this relationship means, since previously we have observed increased urinary clusterin in response to receipt of vancomycin alone. We also found elevations in OPN on days 1-4 for both the vancomycin alone and vancomycin+piperacillin-tazobactam groups; however, OPN is a more general biomarker for acute kidney injury, and not necessarily vancomycin-induced kidney injury ^18^. Based on the results of previous studies as well as these data, KIM-1 appears to be most specific for vancomycin-induced kidney functional changes, with clusterin and OPN being more general markers of kidney injury.

We acknowledge several limitations to this pre-clinical animal study. Baseline (day 0) iohexol GFR measurements were not obtained for these rats. As a result, we were unable to compare post-treatment GFR changes to pre-treatment baseline GFR. However, this model used outbred animals that were all similar at baseline as verified by other biomarkers. In addition, baseline (day 0) urine samples were not obtained for the vancomycin+piperacillin-tazobactam group rats, so baseline CrCL could not be evaluated. Concentrations of urinary biomarkers can be diluted based on urinary volume and multiple approaches have been suggested by investigators. In our study since we were able to collect all urine, we were able to capture the total daily amounts of excreted biomarkers via 24-hour urine samples and compare results over time and between treatment groups. Our findings do not change drastically when data are analyzed as 24-hour concentrations. Finally, the primary analysis for our mixed model did not correct for multiple comparisons. Secondary analyses corrected multiple comparisons using the Bonferroni adjustment. Multiple schools of thought exist in regard to comparing for multiple comparisons ^33^. The primary question in our analysis is: does piperacillin-tazobactam worsen kidney outcomes? By correcting for multiple comparisons, we bias towards type-II errors where no difference is observed in the setting of a true biologic relationship. As such, we have primarily presented data with simple effects of the joint comparisons to controls (i.e. saline) and baseline (day 0 where available) and did not correct for multiple comparisons. As an example in the osteopontin analysis, 15 total comparisons exist and a Bonferroni corrected p-value for significance is p=0.05/15 (or 0.0033 for significance). Importantly, no consistent trends were seen for vancomycin+piperacillin-tazobactam being worse than vancomycin at the less stringent, uncorrected p-value. As would be expected, even less significance is seen in corrected analyses. Rather, vancomycin regularly was worse than vancomycin+piperacillin-tazobactam in uncorrected biomarker analyses and slightly worse in some corrected biomarker analyses. All function studies showed no major worsening of function as measured by iohexol clearance vis-à-vis GFR and creatinine clearance.

## Conclusion

In summary, we have presented the most comprehensive analysis of injury biomarkers and functional kidney measures in the literature to date for rats treated with vancomycin +/− piperacillin-tazobactam (as well as experimental controls). Animals treated with vancomycin alone have worse GFR, injury biomarkers, and histopathology compared to other groups. Our results recapitulate results from previous animal studies and provide further evidence that vancomycin+piperacillin-tazobactam is not associated with additive nephrotoxicity.

## Methods

### Experimental design and animals

The experimental design was similar to previous reports ^15,16,20^. All experiments were conducted at Midwestern University in Downers Grove, Illinois, in compliance with the National Institutes of Health Guide for the Care and Use of Laboratory Animals and were approved under the Institutional Animal Care and Use Committee protocol #3151 ^34^. Male Sprague-Dawley rats (n=24; age: approximately 8 to 10 weeks; mean weight: 275.9 g) were housed in a light- and temperature-controlled room for the duration of the study and allowed free access to water and food. All animals were housed in metabolic cages (Nalgene, Rochester, NY, USA) for 24-hour urine collection, starting on the day prior to study drug dosing (day 0). Animals were assigned to one of four treatment groups in which they received either vancomycin 150 mg/kg/day intravenously over 2 minutes (n=6), piperacillin-tazobactam 1400mg/kg/day intraperitoneal (n=6), vancomycin+piperacillin-tazobactam (n=6), or saline (n=6) for 4 days. Following drug dosing on each day, iohexol 51.8 mg/0.22 mL was administered intravenously over 1 minute. At the end of day 4, all rats were euthanized, and nephrectomies were performed.

### Chemicals and reagents

Rats were administered clinical grade vancomycin hydrochloride (Lot number: 167973; Fresenius Kabi, Lake Zurich, IL, USA), piperacillin-tazobactam sodium (Lot number: 1PU19022; Apollo, Palm Beach Gardens, USA), iohexol [Omnipaque] (Lot number: 15025174; GE Healthcare Inc., Marlborough, MA, USA), and normal saline as a control group (Hospira, Lake Forest, IL, USA). Vancomycin and piperacillin-tazobactam were prepared by weighing and dissolving the powder in purified water (EMD Millipore, Burlington, MA) to achieve final concentrations of 100 mg/mL and 500 mg/mL, respectively. For LCMS analyses, analytical grade iohexol (Lot number: LRAC5648; Sigma-Aldrich, St. Louis, MO), and creatinine (Lot number: LRAB2988; Sigma-Aldrich, St. Louis, MO) were purchased for stock solution preparation; iohexol-d5 (Lot number: LRAC5648; Cayman Chemical, Ann Arbor, MI), and creatinine-d3 (Lot number: 0533175-29; Cayman Chemical, Ann Arbor, MI) were purchased for use as internal standards; LCMS-grade acetonitrile and methanol (VWR, Radnor, PA) were used as mobile phase solvents; formic acid was obtained from Fischer Scientific (Waltham, MA). Frozen, nonimmunized, nonmedicated, pooled plasma (anticoagulated with disodium EDTA) from Sprague Dawley rats (Lot: RAT483100; BioreclamationIVT, Westbury, NY) was utilized for dilution of rat plasma samples.

### Blood and urine sampling

Similar to our previous studies, double jugular vein catheters were surgically implanted 72 hours prior to initiation of the experimental protocol. One catheter was dedicated to blood sample draws, while drug dosing occurred via the other catheter ^12,15,16^. Blood samples were obtained from the catheter at pre-specified timepoints (0, 30, 60, and 240 minutes). Each blood sample (0.2 mL/aliquot) was replaced with the same volume of normal saline (NS) for maintenance of euvolemia. Blood samples were prepared as plasma with EDTA (Sigma-Aldrich Chemical Company, Milwaukee WI, USA) and centrifuged at 3000 g for 10 minutes (Thermo Fisher Scientific, Waltham, MA, USA). Supernatants were collected and frozen at -80℃ until time of batch analysis with liquid chromatography tandem mass spectrometry (LCMS). Urine samples were collected, and volumes were measured every 24 hours, beginning from day 0. Urine samples were centrifuged at 400 g for 5 minutes at 4°C, and the resulting supernatant was collected, aliquoted, and stored at-80℃ until batch analysis of kidney injury biomarkers.

### Removal of kidneys from euthanized rats

Following completion of the dosing protocol, rats were anesthetized with ketamine hydrochloride [Lot number: AH2072X; Covetrus, Portland, ME] and xylazine [Lot number: 0F013; Covetrus, Portland, ME] (100/10 mg/kg) by intraperitoneal injection. Terminal nephrectomies were performed and rats were euthanized by exsanguination through the right atrium while under anesthesia. Left kidneys were formalin fixed for batch histopathological analysis; right kidneys were flash frozen in liquid nitrogen and stored at -80℃.

### Calibration curves in rat plasma

Stock solutions of iohexol and creatinine were prepared at a concentration of 1 mg/mL, while iohexol-d5, and creatinine-d3 were prepared at a concentration of 100 µg/mL. All drugs were dissolved in purified water. An iohexol standard curve was created by diluting stock solution with water to obtain concentrations between 0.5 and 100 µg/mL. Iohexol-d5 was used as the internal standard and was added to each sample to obtain a final concentration of 10 µg/mL. For the creatinine standard curve, stock solution was diluted with water, plasma was then spiked to obtain concentrations between 3.1 and 400 µg/mL. Creatinine-d3 was used as the internal standard and was added to each sample to obtain a final concentration of 10 µg/mL.

For each standard curve concentration, the iohexol or creatinine dilution (4 µL each) was added to 36 µL of blank rat plasma. Iohexol-d5 (4 µL), and creatinine-d3 (4 µL) were added to each standard curve concentration as internal standards. Each standard curve concentration was then mixed with 140 µL of 0.1% formic acid in methanol, vortexed, and centrifuged at 16,000 g for 10 minutes. Resulting supernatant was then transferred to LCMS vials for analysis.

### Sample preparation

For preparation of samples, 4 µL of iohexol-d5, and 4 µL of creatinine-d3 were added to 40 µL of sample rat plasma. Plasma samples at 30 and 60 minutes were diluted 1:10 with blank rat plasma. Each sample was then mixed with 136 µL of 0.1% formic acid in methanol, vortexed, and centrifuged at 16,000 g for 10 minutes. Resulting supernatant was then transferred to LCMS vials for analysis.

### LCMS methods

An Agilent 1260 series liquid chromatography system paired with 6420 triple quadrupole mass spectrometer (Agilent Technologies, Santa Clara, CA) was used to analyze plasma samples. Column temperature was maintained at 20°C. Mobile phase consisted of 0.1% formic acid in water (mobile phase A) and acetonitrile (mobile phase B) for both methods. For analysis of iohexol, a Poroshell 120 analytical column was used (2.7 µm, 100 x 3.0 mm, Part # 695975-302, Agilent Technologies). A gradient method was used to separate the analytes with the following solvent compositions: 5%[A]/95%[B] (0-4 mins), 97%[A]/3%[B] (4.1-7 mins) at a mobile phase flow rate of 0.6 mL/min. Multiple-reaction monitoring mode was used to analyte detection, and the transitions monitored were *m/z* 821.8 to 803.8 and *m/z* 826.8 to 808.8 for iohexol and iohexol-d5, respectively.

Creatinine was analyzed as previously reported ^15^. In brief, a Poroshell 120 analytical column was used (2.7 µm, 50 x 3.0 mm, Part # 699975-302, Agilent Technologies). A gradient method was used to separate the analytes and mobile phase flow rate was 0.6 mL/min. Multiple-reaction monitoring mode was used to analyte detection, and the transitions monitored were *m/z* 114.1 to 44.3 and *m/z* 117.1 to 89.2 for creatinine and creatinine-d3, respectively.

### Determination of urinary biomarkers of AKI

Microsphere-based Luminex xMAP technology was used for the determination of urinary concentrations of KIM-1, clusterin, and OPN, as previously detailed ^16,18,35^. In brief, urine samples were allowed to thaw at ambient room temperature, aliquoted into 96-well plates, and mixed with Milliplex MAP rat kidney toxicity magnetic bead panel 1 (EMD Millipore Corporation, Charles, MO, USA). For each 96-well plate, a separate standard curve was prepared and run, per the manufacturer’s instructions. Results were analyzed and urinary biomarker concentrations were determined using flexible five-parameter (linear and logarithmic scale) curve-fitting models (Milliplex Analyst 5.1; VigeneTech, Carlisle, MA, USA).

### GFR estimation

GFR was assessed by iohexol clearance as described in model building below. Iohexol was administered intravenously as an undiluted solution on each dosing day, after administration of the study drug treatment. Rats received once-daily doses of iohexol 51.8mg/0.22 mL, given over 1 minute, on experimental days 1 through 4.

### Creatinine clearance (CrCL) calculation

Urinary and plasma creatinine concentrations were quantified by LCMS. CrCL was calculated according to the equation: CrCL = (Urine _Creatinine concentration_ x Urine _Volume_) / (Serum _Creatinine concentration_ x Time _minutes_) ^36^. The resulting CrCL was then normalized to a rate in mL/min/100 g body weight, according to each individual rat’s daily weight.

### Histopathological analysis of kidneys

Formalin-fixed kidneys were sent for batch histopathological analysis (IDEXX Bioanalytics, Columbia, MO, USA). Similar to our previous studies, blinded samples were scored according to the Predictive Safety Testing Consortium (PSTC) semi-quantitative grading system by a trained pathologist ^16,35^. Scores ranged from 0 to 5: 0 indicating no abnormalities, 1 indicating minimal, very few, or very small abnormalities, 2 indicating mild, few, or small abnormalities, 3 indicating moderate, moderate amount, or moderate sized abnormalities, 4 indicating marked, many, or large abnormalities, and 5 indicating massive, extended number, or very large abnormalities noted. We utilized the worst overall score, representing total gross kidney damage, and worst tubular scores in our evaluation for end-organ damage.

### Model building

In order to describe iohexol clearance, a two-compartment model was created (Monolix 2021R1; Lixoft, Antony, France). Fixed estimates for peripheral compartment volume (V2) and intercompartmental transfer (Q) were derived from the base two-compartment model fitting. Random effects were estimated for clearance (CL) and central volume (V1). Evaluated covariates included log-transformed weight and treatment group. To capture daily clearance changes, each experimental day was considered a separate occasion with clearance that could vary. Selection of the final model was based on the Akaike Information Criterion (AIC), between subject variability of the population estimates, goodness-of-fit plots for observed versus predicted, and the rule of parsimony ^37^. Lowest AIC was utilized for any discrepancies. Empirical Bayes Estimates (i.e. individual Bayesian posteriors) were utilized for individual animal parameters.

### Statistical analysis

A mixed-effects, restricted maximum likelihood estimation regression was used for comparison of urine output, mean weight loss, GFR, CrCL, and urinary biomarkers among the treatment groups, with repeated measures occurring over days; measures were repeated for each individual rat (STATA version 16.1, StataCorp LLC, College Station, TX, USA) ^16^. Margins were calculated for a full factorial of the variables, i.e. main effects for each variable and interactions. Referent groups were pre-treatment baseline values (for all outcomes except GFR in which day 1 was the referent group) and saline as a treatment. The primary analysis reports the simple effects from joint tests of drug treatment group within each level of treatment day. To avoid biasing toward the null (i.e.piperacillin-tazobactam does not change kidney outcomes), simple effects were the primary comparison. However, highly conservative Bonferroni corrected p-values are additionally reported. In addition, we indicate which interactions of treatment group by time are significant at a P<0.05 with the notation ** in the results and Supplemental table 1. Spearman’s rank correlation coefficient with a Bonferroni correction was used to assess correlations between kidney injury biomarker expression (e.g. KIM-1, clusterin, OPN) and function (e.g. GFR) by treatment day. Logarithmic transformations of data were considered as necessary for normalization of data distribution. Locally weighted scatterplot smoothing (LOWESS) trendlines with 95% confidence intervals are provided for graphical output to remain agnostic to outcome variable relationship over time (and visually allow comparisons between treatment groups). All utilized tests were two-tailed, with an *a priori* level of statistical significance set at α = 0.05.

## Acknowledgements

MS gratefully acknowledges funding from the National Institute of Allergy and Infectious Diseases under award number R21-AI149026. The content is solely the responsibility of the authors and does not necessarily represent the official views of the National Institutes of Health

## Author contributions

JC had primary responsibility for sample collection, analytical quantification, data analysis, and authored the paper. GP performed all animal surgeries. SM, KV, and EL assisted with care of animals and sample collection. EB assisted with data analysis. MS designed and oversaw experimental conduct, including data analysis. All authors discussed and commented on the manuscript.

## Data availability

Data that support the findings of this study are available from the corresponding author upon reasonable request. Data sharing will be subject to standard Data Use Agreements from Midwestern University.

## Animal welfare statement

Care and treatment of the experimental animals was in accordance with local and international guidelines on the ethical use of laboratory animals. All procedures used in this study were approved by the Midwestern University Institutional Animal Care and Use Committee under protocol #3151 and conducted in accordance with the ARRIVE guidelines.

